# GABA produced by multiple bone marrow cell types regulates hematopoietic stem and progenitor cells

**DOI:** 10.1101/2025.04.30.651482

**Authors:** Cesi Deng (邓策思), Adedamola Elujoba-Bridenstine, Ai Tang Song, Rylie M. Ceplina, Casey J. Ostheimer, Molly C. Pellitteri Hahn, Cameron O. Scarlett, Owen J. Tamplin

## Abstract

Hematopoietic stem and progenitor cells (HSPCs) maintain homeostasis of the blood system by balancing proliferation and differentiation. Many extrinsic signals in the bone marrow (BM) microenvironment that regulate this balance are still unknown. We report gamma aminobutyric acid (GABA) metabolite produced in the BM as a regulator of HSPCs. Deletion of the glutamate decarboxylase enzymes (Gad1/2) that produce GABA in either B lineages or endothelial cells (ECs) leads to a moderate reduction in BM GABA levels and HSPC number, suggesting both cell types are GABA sources. However, simultaneous blockade of GABA production from both hematopoietic cells and ECs resulted in a greater reduction of both GABA levels and HSPC numbers. Lower GABA levels in the BM altered the gene expression profile of HSPCs, with expression reduced for proliferation-associated genes and increased for B lineage genes. Our findings suggest GABA from multiple sources coordinates to regulate HSPC activity.

**Highlights:**

- GABA is produced by B cells and endothelial cells in the bone marrow
- Lower GABA level in the bone marrow reduces HSPC proliferation
- Lower GABA level primes HSPCs to upregulate B cell differentiation programs

**eTOC blurb:** Tamplin and colleagues functionally test production of GABA metabolite in the bone marrow microenvironment as a regulator of hematopoietic stem and progenitor cells. Conditional deletion of GAD enzymes in B cells and endothelial cells demonstrated both are sources of GABA. Lower GABA level primed HSPCs to reduce proliferation and upregulate B cell differentiation programs.

**Graphical Abstract:** 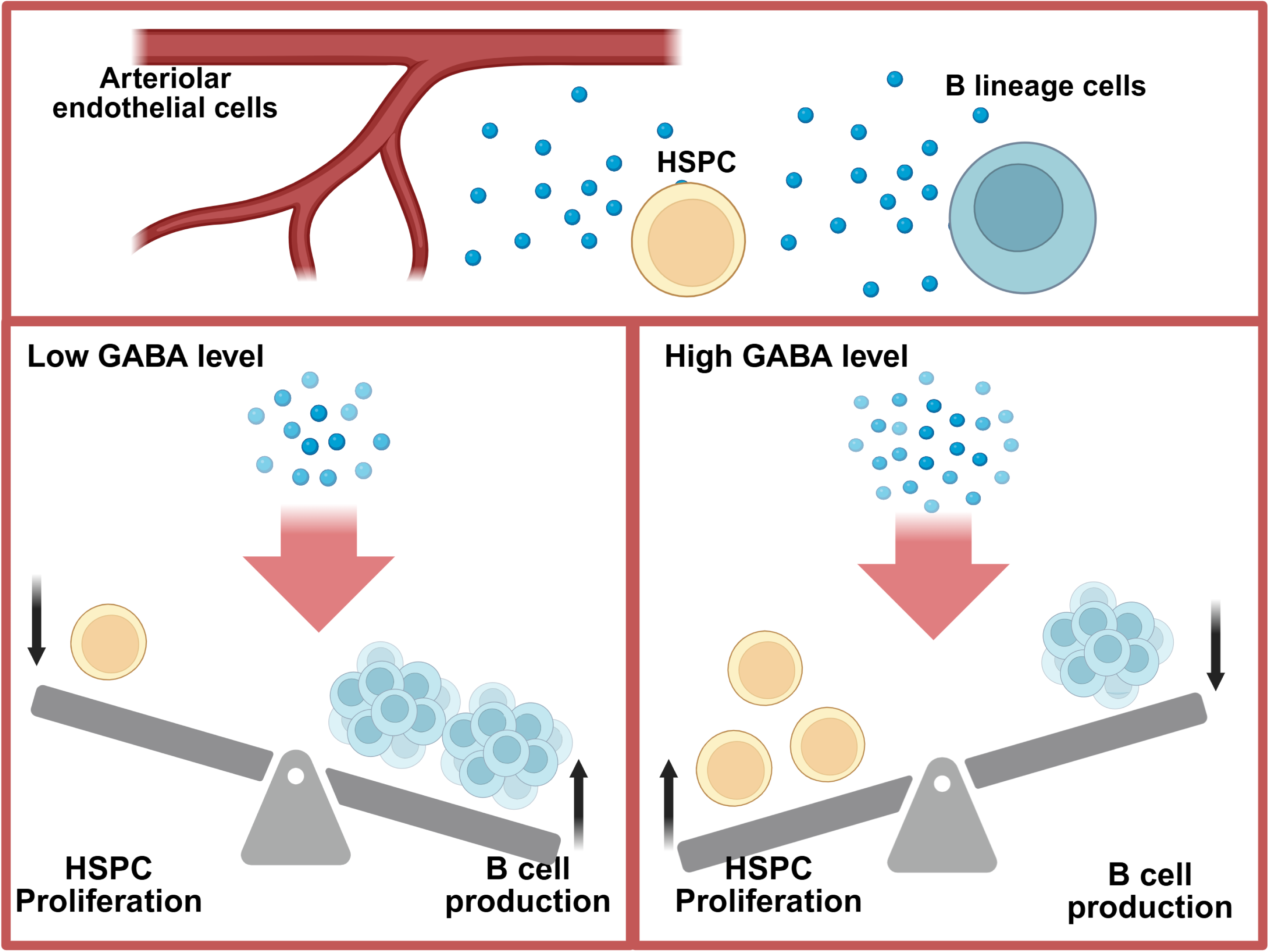

## Introduction

Hematopoietic stem and progenitor cells (HSPCs) replenish the entire blood system throughout life (Pinho and Frenette, 2019). The balance between HSPC proliferation and differentiation is tightly regulated by extrinsic signals in their microenvironment, also known as the niche. In adults, bone marrow (BM) is the primary site of hematopoiesis and is made up of a highly complex array of different niche cell types. Although research has focused on the cytokines, other growth factors, receptor ligands, and adhesion molecules produced by these different niche cells, there are also understudied factors such as metabolites and neurotransmitters that play a role in regulating HSPC activity. For example, peripheral nerves can provide adrenergic signals that act on stromal cells to regulate HSPC mobilization (Méndez-Ferrer et al., 2008), and dopamine can act directly acting on neuroreceptors on the HSPC surface (Liu et al., 2021). The full range of secreted immune metabolites are reviewed elsewhere (Zhang et al., 2022).

Gamma amino-butyric acid (GABA) is a major neurotransmitter in the central nervous system. GABA binds to two classes of receptors, ionotropic GABA_A_ receptor and metabotropic GABA_B_ receptor. Interestingly, both types of GABA receptors have been found on HSPCs. GABA type A receptor subunit rho 1 (GABRR1) marks a subset of HSPCs and its deletion inhibited megakaryocyte differentiation and reduced blood platelets (Zhu et al., 2019). In our previous work, we showed that loss of GABA type B receptor subunit 1 (GABBR1) reduced proliferation and reconstitution capacity of hematopoietic stem cells (HSCs), and impaired B cell production (Shao et al., 2021). While these findings suggest GABA has a pleiotropic role in regulating HSPC activity, the sources of GABA in the BM remain unclear. We examined the spatial distribution of GABA in the BM using imaging mass spectrometry and found it was enriched in the endosteal region of the mouse femur (Shao *et al*., 2021). GABA production is catalyzed by glutamate decarboxylases 1 and 2 (GAD1/2) (Erlander et al., 1991). Work by us and others showed *GAD1* but not *GAD2* is expressed in human and mouse B cells that are also enriched for GABA metabolite (Shao et al., 2021, Zhang et al. 2021). We sought to block GABA production from various BM niche cell types. We show that B cells and endothelial cells produce GABA, which is required for the maintenance of the HSPC pool. HSPCs are primed to commit to the B cell lineage when GABA level in the microenvironment is low. We propose that GABA serves as an environmental cue to regulate the decision between HSPC proliferation and B cell production to maintain homeostasis of the B cell pool.

## Results

### B cells are a source of GABA in the bone marrow niche

To determine if B cells are an autonomous source of GABA, we tracked GABA production throughout B cell development using the OP9 stromal cell co-culture system (Holmes and Zuniga-Pflucker, 2009; Shao *et al*., 2021). In brief, HSPCs (Lin-/c-Kit+/Sca-1+) sorted from wild-type (WT) mice were co-cultured with stromal cells and examined at 3-day intervals. GABA level in the co-culture media was determined by mass spectrometric analysis, and B lineage cell production was determined by flow cytometry (Fig. 1A-B). We found Cd19+ cells emerged on Day 9 and underwent rapid expansion in the following three days (Fig. 1A-C). Our baseline was GABA level in media and OP9 cells alone that do not produce GABA, to normalize against the small amount of GABA present in fetal bovine serum. During B cell differentiation the GABA level remained low until Day 9 and then increased dramatically at Day 12 (Fig. 1C). The synchronization of B lineage cell emergence and GABA production demonstrated B cells are a source of GABA. This autonomous production of GABA by B cells during differentiation was consistent with the results of others that detected GABA in B cells purified from human peripheral blood and mouse lymph nodes (Zhang et al., 2021).

**Figure 1.**
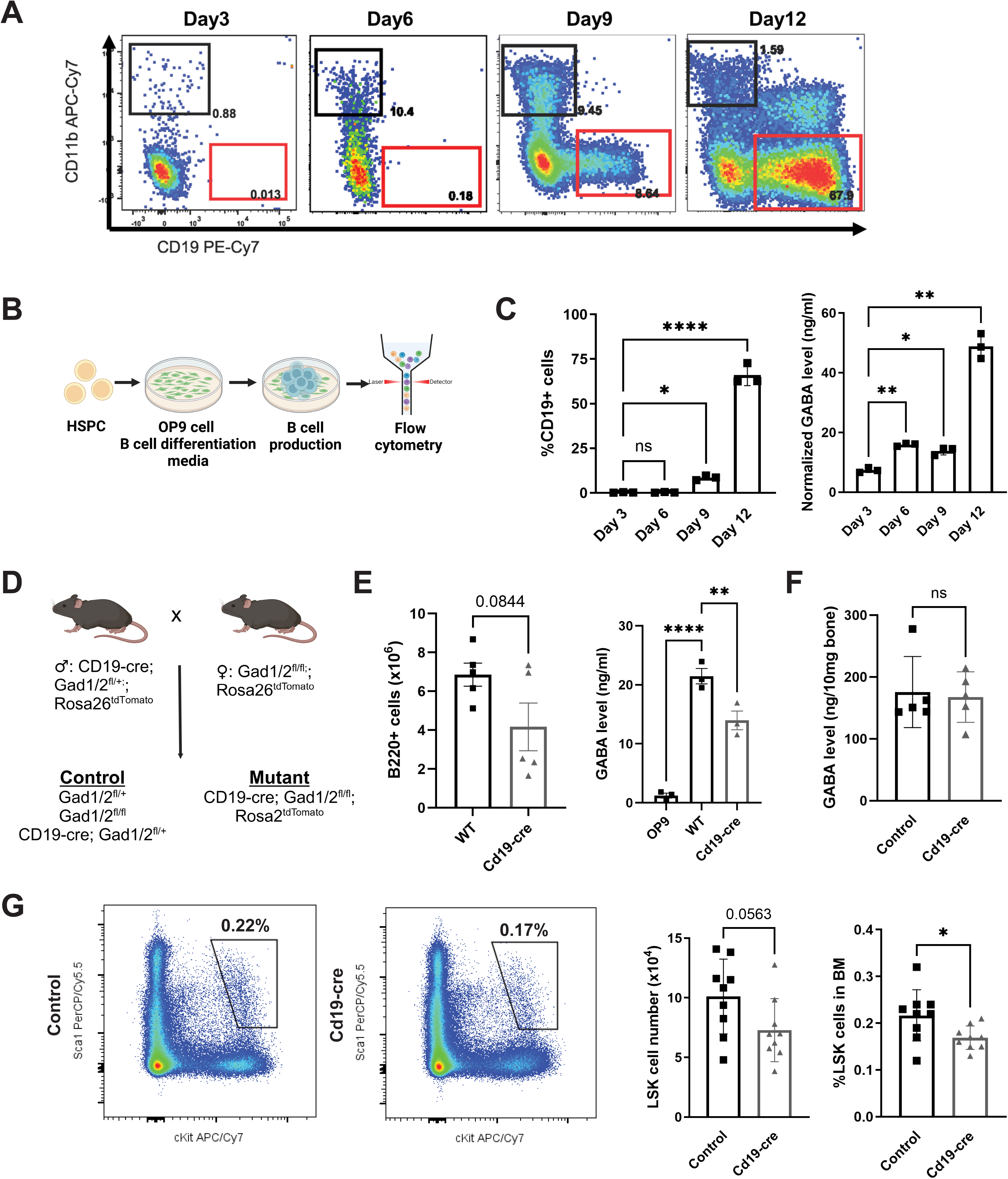
B cells are a source of GABA in the bone marrow. (A) Representative flow plots showing the emergence of B lineage cells in the OP9 co-culture. (B) Schematic of HSPC OP9 B cell differentiation co-culture. (C) Left: Quantification of the percentage of Cd19+ B cells at different timepoints; Right: mass spectrometric quantification of GABA level in the media throughout B cell differentiation. (D) Breeding scheme to generate *Cd19-Cre;Gad1/2^fl/fl^;Rosa26^tdTomato^* mice and control mice. (E) Left: Quantification of B cells produced during differentiation of HSPCs; Right: GABA level in the media from *Cd19-cre;Gad1/2^fl/fl^;Rosa26^tdTomato^*mice and WT mice at Day 12. (F) Mass spectrometric quantification of GABA in *Cd19-cre;Gad1/2^fl/fl^;Rosa26^tdTomato^*and control mouse femur (n = 5 mutants vs n = 5 control). (G) Left: Representative flow plots showing the HSPC gates (Lin-/Sca-1+/c-Kit+); Right: Quantification of LSK cell abundance in *Cd19-cre;Gad1/2^fl/fl^;Rosa26^tdTomato^* mice and control mice (n = 9 mutants vs n = 9 control). Data is represented as mean ± SEM (*p<0.05, **p<0.01, ***p<0.001, ****p<0.0001), Welch’s test and one-way ANOVA with Tukey’s test were performed.

To confirm the *in vivo* functional requirement of GABA production by B cells in the BM niche, we generated mouse models with B cell-specific deletion of *Gad1/2*. We crossed *Cd19-Cre* (Rickert et al., 1997) and *Gad1/2^fl/fl^;Rosa26^tdTomato^*mouse lines (Meng et al., 2016) to generate *Cd19-cre;Gad1/2^fl/fl^;Rosa26^tdTomato^*(Fig. 1D). *Cd19-Cre* activity was shown by the percentage of double positive Cd19+/tdTomato+ cells (Fig. S1A). We isolated HSPCs from *Cd19-cre;Gad1/2^fl/fl^;Rosa26^tdTomato^* mice and induced their differentiation into B cells using the OP9 co-culture system. There was significant reduction of GABA level in co-cultures seeded with HSPCs from *Cd19-cre;Gad1/2^fl/fl^;Rosa26^tdTomato^*mice (Fig. 1E).

We used mass spectrometry to measure GABA levels in femurs from *Cd19-Cre;Gad1/2^fl/fl^;Rosa26^tdTomato^* mice but did not find a significant difference compared to controls (Fig. 1F). Consistent with previously published results that used the B lineage-specific conditional *Gad1* knockout model *Mb1-Cre;Gad1^fl/fl^* (Zhang *et al*., 2021), we also observed no change in BM HSPC numbers, number of BM B220+ cells, or percentage of BM and peripheral blood (PB) B220+ cells (Fig. S2A-C). Interestingly, we did observe a decrease in the proportion of BM HSPCs (Fig. 1G), a phenotype that was not tested in the *Mb1-Cre;Gad1^fl/fl^* model (Zhang *et al*., 2021), suggesting B cell-derived GABA may directly regulate HSPCs.

### Endothelial cells are a source of GABA in the bone marrow niche

Data from our group and others found BM endothelial cells (BMECs) express *Gad1* and are therefore a potential source of GABA (Shao *et al*., 2021; Zhu *et al*., 2019). Notably, published mouse BM stroma single cell RNA-seq datasets do not show *Gad1* expression in any populations (Baryawno et al., 2019), while quantitative PCR and bulk RNA-seq detect its expression in B cells and BMECs (Shao *et al*., 2021; Zhang *et al*., 2021; Zhu *et al*., 2019), suggesting *Gad1* transcript may be expressed at low levels. To functionally test if ECs contribute to GABA production in BM, we generated an endothelial-specific *Gad1/2* deletion by crossing *Cdh5-CreERT2* (Sörensen et al., 2009) and *Gad1/2^fl/fl^;Rosa26^tdTomato^* lines to generate *Cdh5-CreERT2;Gad1/2^fl/fl^;Rosa26^tdTomato^* mice (Fig. 2A). Tamoxifen was administered at 5-6 weeks and experiments were performed at 8-12 weeks. *Cdh5-CreERT2* activity was confirmed by the presence of tdTomato+ cells in the BM (Fig. S1B). There was no significant decrease in GABA levels in *CreERT2;Gad1/2^fl/fl^;Rosa26^tdTomato^* BM (Fig. 2B); however, there was a reduction in both number and proportion of HSPCs (Fig. 2C). No change was detected in BM cellularity, CLPs or B cells, or PB myeloid or B cells (Fig. S2D-H). These results demonstrated that *Gad1/2* deletion from BMECs did not broadly impact BM hematopoiesis or GABA levels, however it was sufficient to modulate the proportion and number of HSPCs.

**Figure 2.**
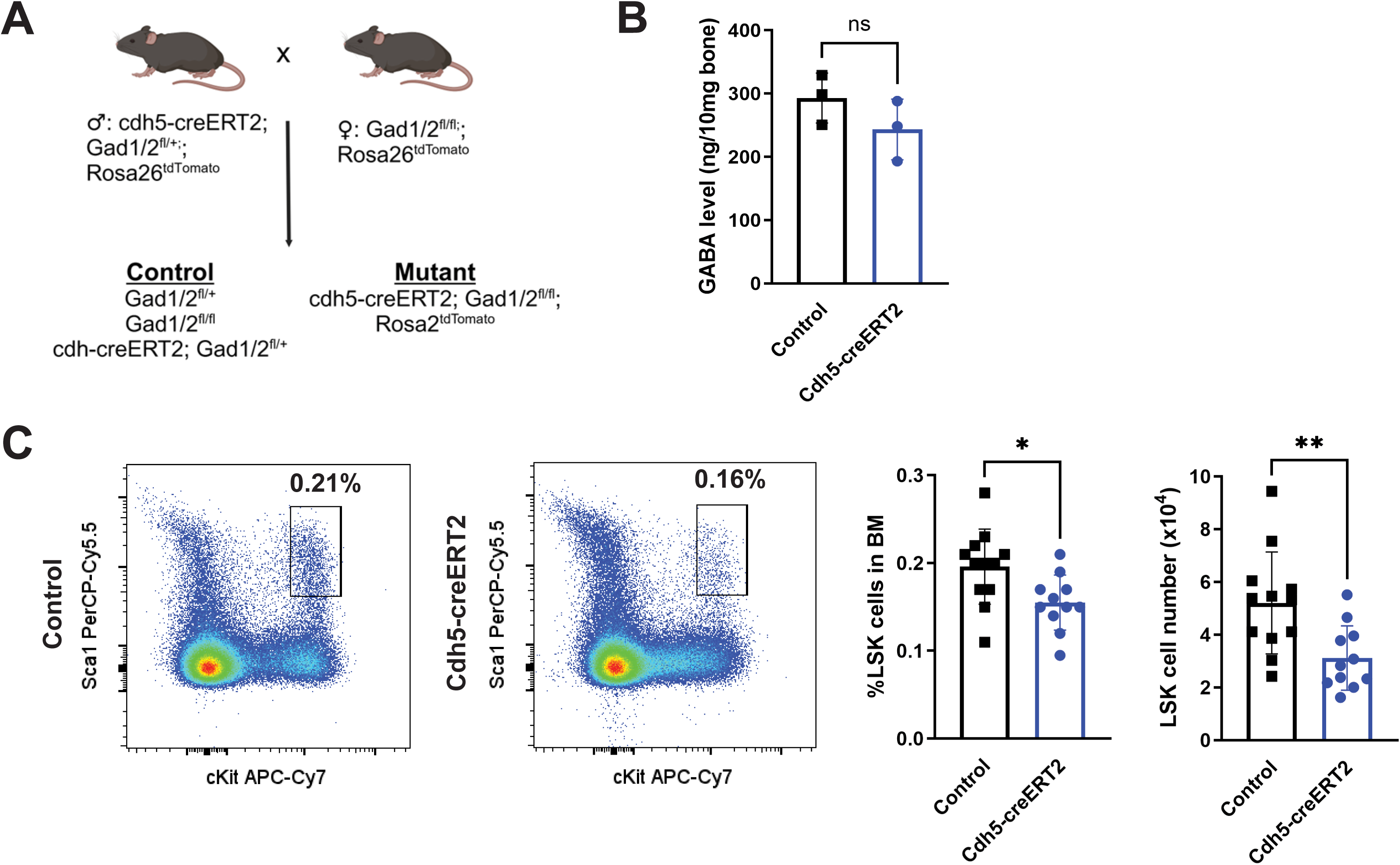
Bone marrow endothelial cells are a source of GABA in the niche. (A) Breeding scheme to generate *Cdh5-CreERT2;Gad1/2^fl/fl^;Rosa26^tdTomato^*mice and control mice. (B) Mass spectrometric quantification of GABA in *Cdh5-CreERT2;Gad1/2^fl/fl^;Rosa26^tdTomato^* and control mouse femur (n = 3 mutants vs n = 3 control). (C) Left: Representative flow plots showing the HSPC gates (Lin-/Sca-1+/c-Kit+); Right: Quantification of LSK cell abundance in *Cdh5-CreERT2;Gad1/2^fl/fl^;Rosa26^tdTomato^* mice and control mice (n = 11 mutants vs n = 12 control). Data is represented as mean ± SEM (*p<0.05, **p<0.01), Welch’s test was performed.

#### Hematopoietic and BMEC-derived GABA coordinates to regulate HSPC activity

To understand the contribution of multiple sources of GABA in the BM, we generated a *Tie2-Cre;Gad1/2^fl/fl^;Rosa26^tdTomato^*mouse model (Fig. 3A). *Tie2-Cre* is expressed in all ECs during embryonic development and into adulthood (Kisanuki et al., 2001). In the embryo, the hemogenic endothelium gives rise to all definitive HSPCs, meaning in the adult *Tie2-cre* model all ECs and hematopoietic cells have undergone Cre excision (Tang et al., 2010). Therefore, the *Tie2-Cre;Gad1/2^fl/fl^;Rosa26^tdTomato^* mouse model enabled us to evaluate the effect of *Gad1/2* deletion in all ECs and hematopoietic cells, inclusive of both B cells and BMECs. Cre activity was confirmed by detection of tdTomato+ cells in the BM (Fig. S1C). We measured GABA levels in the BM of *Tie2-Cre;Gad1/2^fl/fl^;Rosa26^tdTomato^* mice and detected a ∼60% reduction compared to control mice (Fig. 3B). Considering that deletion of *Gad1/2* from B cells (Fig. 1F) or ECs alone (Fig. 2B) did not lead to a significant reduction of BM GABA, our results using the *Tie2-Cre* model suggest B cells and BMECs synergize to sustain homeostatic BM GABA levels. As in the EC-specific *Cdh5-CreERT2;Gad1/2^fl/fl^;Rosa26^tdTomato^* model (Fig. 2C), we found a decrease in HSPC percentage and number in *Tie2-Cre;Gad1/2^fl/fl^;Rosa26^tdTomato^*BM (Fig. 3C). We did not observe any changes in BM cellularity, CLPs, or B220+ cells, or PB myeloid cells (Fig. S2I-L). We did observe a slight reduction in the percentage of PB B220+ cells (Fig. 3D). Although the *Cd19-Cre*, *Cdh5-CreERT2*, and *Tie2-Cre* models all showed a decrease in the proportion of BM HSPCs, only the combined hematopoietic and EC-specific deletion of *Gad1/2* with *Tie2-Cre* resulted in a significant reduction in BM GABA levels. This shows that GABA is produced by multiple sources at steady state to regulate HSPC activity in the BM niche.

**Figure 3.**
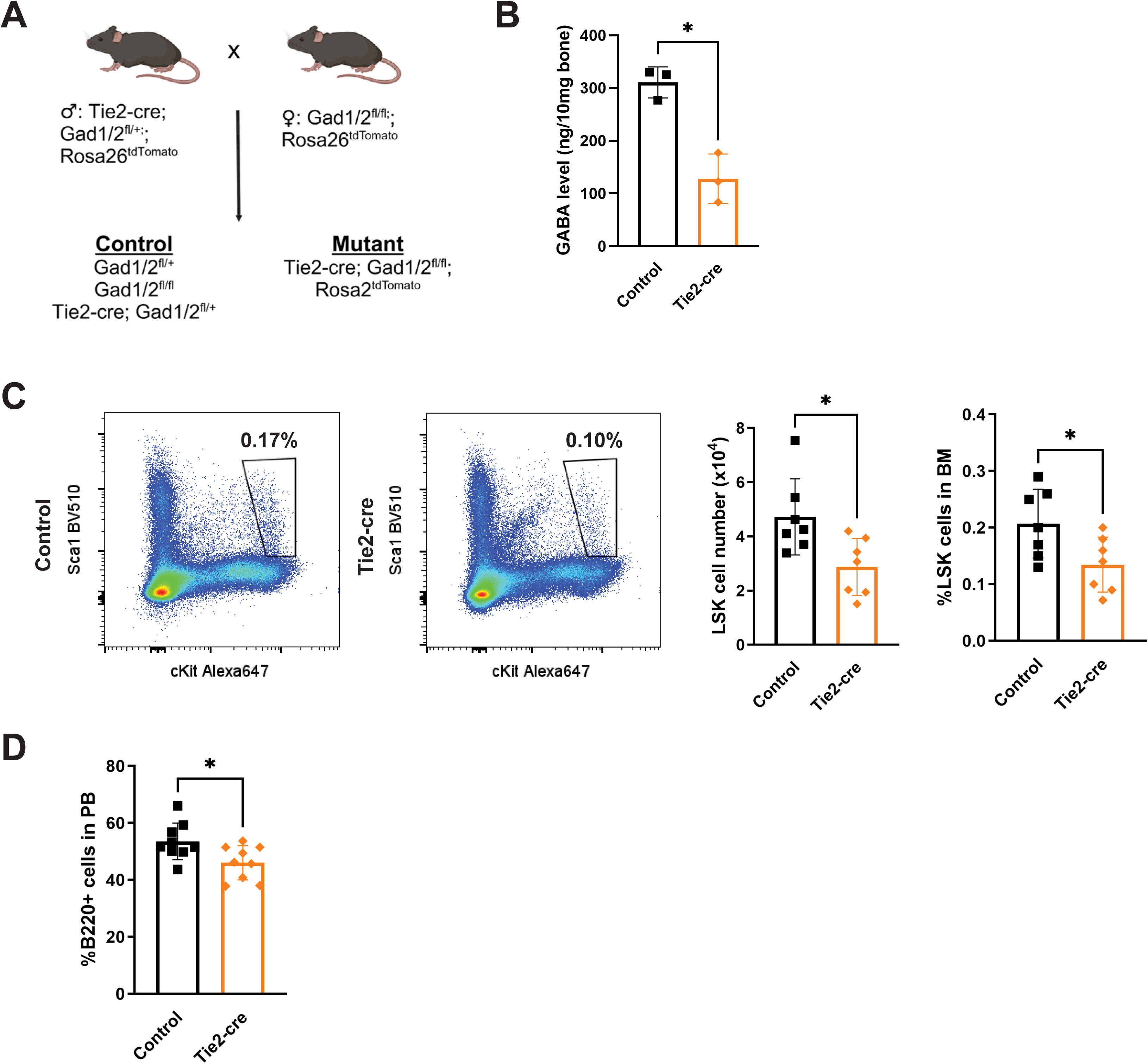
BMEC and hematopoietic cell-derived GABA synergizes to regulate HSPC activity. (A) Breeding scheme to generate *Tie2-Cre;Gad1/2^fl/fl^;Rosa26^tdTomato^* mice and control mice. (B) Mass spectrometric quantification of GABA in *Tie2-Cre;Gad1/2^fl/fl^;Rosa26^tdTomato^*and control mouse femur (n = 3 mutants vs n = 3 control). (C) Left: Representative flow plots showing the HSPC gates (Lin-/Sca-1+/c-Kit+); Right: Quantification of LSK cell abundance in *Tie2-Cre;Gad1/2^fl/fl^;Rosa26^tdTomato^* mice and control mice (n = 7 mutants vs n = 7 control). (D) Percentage of B220+ cells in the peripheral blood in *Tie2-Cre;Gad1/2^fl/fl^;Rosa26^tdTomato^*mice and control mice (n = 9 mutants vs n = 9 controls). Data is represented as mean ± SEM (*p<0.05), Welch’s test was performed.

#### Reduced BM GABA levels induced a shift in gene expression profiles of HSPCs

To determine if attenuated BM GABA levels result in intrinsic molecular changes in HSPCs, we isolated HSPCs from *Cd19-Cre*, *Cdh5-CreERT2*, and *Tie2-Cre;Gad1/2^fl/fl^;Rosa26^tdTomato^*models and performed bulk RNA-sequencing. Among the total 21,114 genes detected, there were differentially expressed genes (DEGs) in HSPCs of each model when compared to WT: 494 in *Cd19-Cre* (367 upregulated, 127 downregulated); 284 in *Cdh5-CreERT2* (131 upregulated, 153 downregulated); 200 in *Tie2-Cre* (73 upregulated, 127 downregulated; FDR < 0.1, |log2FC| > 1; Fig. 4A-B, Fig. S3A). Comparison of DEGs in each mutant showed 21 genes were commonly upregulated upon GABA level reduction, including HSC genes *Hlf*, *Emcn*, and *Fgd5* (Fig. 4A-C) (Calvanese et al., 2022; Engelhard et al., 2024; Gazit et al., 2014; Holmfeldt et al., 2016; Matsubara et al., 2005; Reckzeh et al., 2018; Ueno et al., 2001). Together, this demonstrates that HSPCs respond to changes in GABA produced by BMECs and hematopoietic cells.

**Figure 4.**
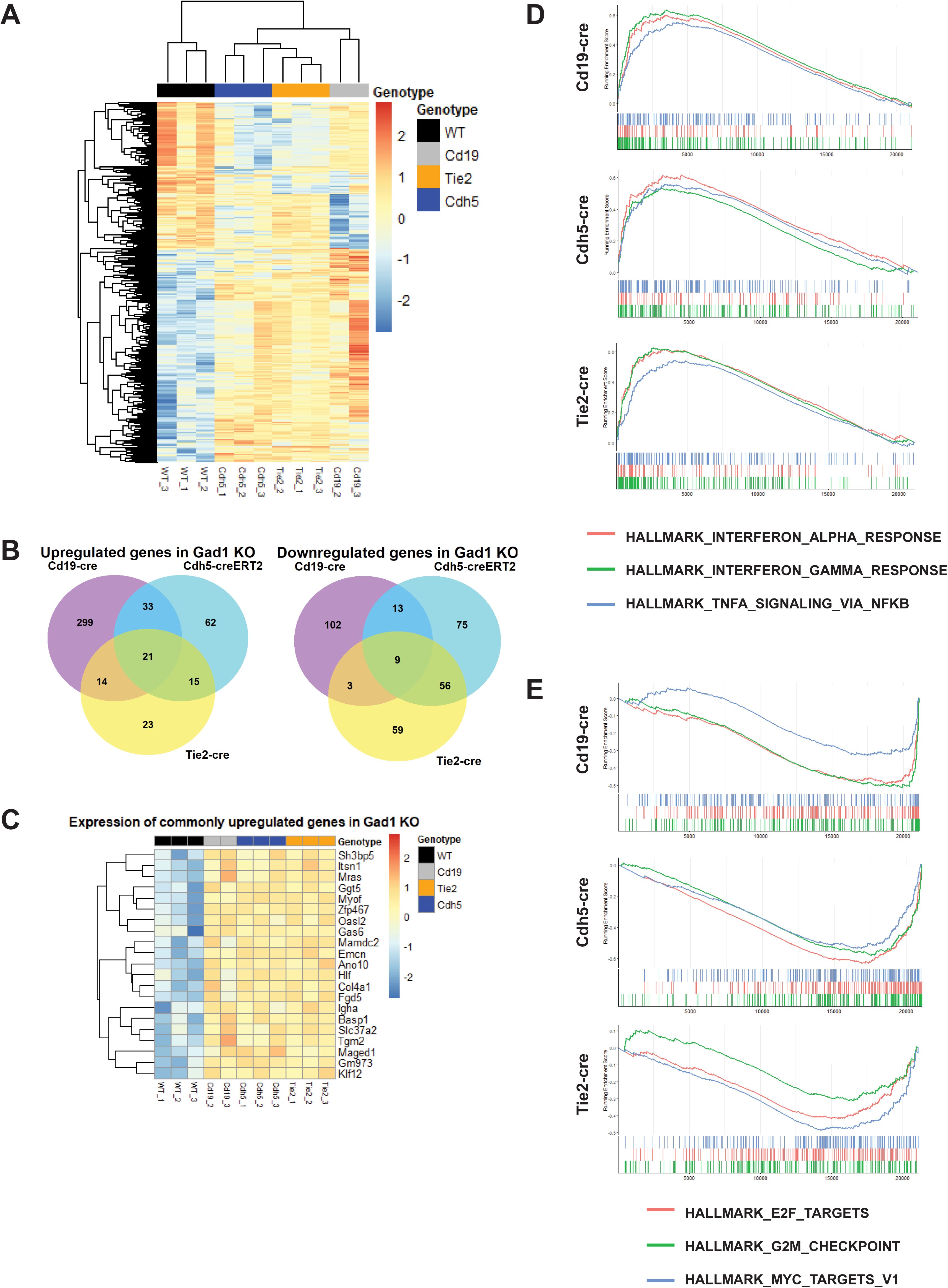
Reduction in GABA level induced transcriptional changes in HSPCs. (A) Heatmap of hierarchical clustering based on differentially expressed genes in *Gad1/2^fl/fl^* mutant models, (|logFC| >1, FDR < 0.1). (B) Venn diagram of genes that were commonly or differentially expressed in *Gad1/2^fl/fl^* mutant models. (C) Heatmap of the expression level of 21 genes that were commonly upregulated in *Gad1/2^fl/fl^* mutant models. (D) GSEA enrichment plots showing upregulated pathways in *Gad1/2^fl/fl^* mutant models. (E) GSEA enrichment plots showing downregulated pathways in *Gad1/2^fl/fl^* mutant models.

Our previous analysis of GABA receptor *Gabbr1* loss-of-function mutant HSPCs found they were less proliferative, less responsive to poly(I:C), and had defective B cell differentiation (Shao *et al*., 2021). Intriguingly, phenotypic *Gabbr1* null HSPCs exhibited a shift in transcriptional profiles from undifferentiated progenitor to more B lineage committed progenitor (Shao *et al*., 2021). We hypothesized loss of *Gabbr1* in HSPCs would simulate attenuated GABA signaling and therefore may mimic HSPCs from a low GABA microenvironment. To test this, we considered DEGs separately in the *Cd19-Cre*, *Cdh5-CreERT2*, and *Tie2-Cre* models and performed gene set enrichment analysis (GSEA) for HSPCs from each model (Fig. 4D,E; Fig. S3C-E; Supplementary Tables). Like *Gabbr1* null HSPCs, we found B cell receptor signaling pathway and B cell activation were enriched in all compared to WT (Fig. S3C-E). We also found interferon signaling pathways were highly upregulated in all *Gad1/2^fl/fl^* models (Fig 4D). This is consistent with our previous finding that showed a positive correlation between *Gabbr1* expression in HSPCs and the expression of genes related to interferon signaling (Shao *et al*., 2021), further supporting the GABA-GABBR1 axis in regulating HSPC differentiation and immune response. Consistent with reduced proliferative capacity, E2F and MYC targets, and G2M checkpoint genes were under-represented in *Gad1/2^fl/fl^* models compared to WT (Fig 4E). Together, this showed that phenotypic HSPCs from a low GABA microenvironment have a similar intrinsic profile to HSPCs that lack GABBR1 receptor signaling.

There were many similarities between *CD19-Cre*, *Cdh5-CreERT2*, and *Tie2-Cre* models, however we wanted to consider if there were differences in HSPC regulation that were dependent on the cellular source of GABA. The hierarchical clustering of DEGs from all models showed clusters that were unique to B lineages, BMECs, or both (Fig. 4A). Intriguingly, a curated list of B lineage genes (Miyai et al., 2018) found some of these markers were only strongly upregulated in the *Cd19-Cre* model (e.g., *Cd79a, Cd79b, Vpreb1*; Fig. S3B; Supplementary Tables). *Mecom*, an important regulator of HSCs (Voit et al., 2023), was only upregulated in models that deleted *Gad1/2* from BMECs (i.e., *Cdh5-CreERT2 and Tie2-Cre*). These findings suggest that HSPCs may be differentially regulated by GABA depending on their location in the BM niche, and their proximity to B cells or BMECs. Together, our data shows reduced production of GABA in the BM niche promotes HSPC priming towards B cell lineage commitment, while suppressing HSPC proliferation. Overall, our results reveal a role for GABA as a molecular switch to regulate HSPC activity and B lineage priming.

## Discussion

GABA is well-known as an inhibitory neurotransmitter in the central nervous system, but it has also been detected in peripheral organs, including the peripheral nervous system, gastrointestinal track, pancreas, and liver (Zhang *et al*., 2022). In these peripheral tissues, GABA has diverse functions beyond transmitting neural signals, such as regulating immune cell activity in lymph nodes. We demonstrate that GABA serves as an environmental cue in the BM to regulate HSPC differentiation into B cells and maintain the stem cell pool. Despite detectable GABA in multiple peripheral tissues, the sources of GABA remain largely unknown. We show here that B cells and BMECs synergize to maintain homeostatic levels of GABA. Interestingly, GABA level in the BM was not fully reduced even when *Gad1/2* was deleted from both hematopoietic cells and ECs using the *Tie2-Cre* model, suggesting other unknown GABA sources. A survey of published mouse BM stromal cell single cell RNA-seq datasets did not identify any cells that express *Gad1* or *Gad2* (Baryawno *et al*., 2019), possibly because mRNA expression levels are below the limit of detection using this method. While GAD1 is considered the primary pathway for GABA biosynthesis, other mechanisms have been found, including catalysis of putrescine by aldehyde dehydrogenases (Kim et al., 2015; Li et al., 2025). Further investigation will be needed to identify all the sources of GABA production in the BM microenvironment.

HSPCs sorted from the BM of *CD19-Cre*, *Cdh5-CreERT2*, and *Tie2-Cre* models shared many phenotypic similarities. This included an upregulation of immune response genes, including immunoglobulin genes and B cell blueprint genes. This observation was consistent with phenotypic HSPCs sorted from *Gabbr1* null mutant BM, where later-stage B cell commitment genes and pro-B cell-signature genes were highly upregulated (Shao *et al*., 2021). Strikingly, HSPCs from our *Gad1/2*-deficient BM niche models also had downregulation of genes associated with proliferation, another hallmark of *Gabbr1* null mutant HSPCs. These shared changes in HSPCs that lack the GABBR1 receptor, or reside in BM with lower GABA levels, suggests the GABA-GABBR1 axis is important for balancing a decision between HSPC proliferation and differentiation, specifically towards B lineages.

Although we observed many similarities in HSPCs from different *Gad1/2*-deficient BM niche models, there were differences as well, such as B lineage-specific genes *Cd79a*, *Cd79b*, and *Vpreb1* that were uniquely upregulated in *Cd19-cre^+^;Gad1/2^fl/fl^* mutant HSPCs. These transcriptional differences likely arise from the blockade of GABA production in different niche cell types. Spatial mapping of lymphopoiesis in the sternum has revealed that common lymphoid progenitors (CLPs) reside in an arteriolar niche, surrounded by Pre-Pro-B, Pro-B and Pre-B cells (Wu et al., 2024). Within this B cell production line, more lineage-committed cells are located further away from CLPs, suggesting spatial heterogeneity in GABA production. While both endothelial cells and B lineage cells are GABA sources, lymphoid progenitors are proximal to arterioles and more distant from Pro-B and Pre-B cells (Wu *et al*., 2024). Therefore, the blockade of GABA production from either or both sources could differentially impact HSPC activity depending on their spatial relationships. Understanding the spatial heterogeneity of niche factors will provide more insights into how specialized niches support hematopoiesis.

Downstream differentiated hematopoietic cells have been recognized as regulators of their own upstream HSPCs through feedback mechanisms (Kirouac et al., 2010). For example, platelet-biased vWF + HSCs associate with megakaryocyte niches and are regulated by CXCL4 produced by megakaryocytes (Pinho et al., 2018). We demonstrate here that GABA-producing B cells regulate GABBR1+ HSPCs, emphasizing the importance of feedback mechanisms in the BM niche to maintain homeostasis. Pharmacological modulation of these mechanisms could lead to repurposing of clinical neuromodulators to target the hematopoietic and immune systems for therapeutic benefits.

## Methods

### Mice

Cd19-Cre (JAX stock #006785) (Rickert *et al*., 1997), *Tie2-Cre* (JAX stock #004128) (Koni et al., 2001), *Gad1^lox^::Gad2^lox^::Rosa26^tdTomato^*(JAX stock #031800) (Meng *et al*., 2016), and C57BL/6 mice were purchased from the Jackson Laboratory. *Cdh5-CreERT2* was generated by Ralf Adam’s group (Sörensen *et al*., 2009). To generate *Gad1/2* deletion in different cell types, *Cd19-Cre*, *Cdh5-CreERT2*, and *Tie2-Cre* were bred to *Gad1^lox^::Gad2^lox^::Rosa26^tdTomato^*. Genotyping was performed by Transnetyx. At 5-6 weeks of age, *Cdh5-CreERT2;Gad1/2^fl/fl^* mice or Cre-negative littermate controls were injected with 1.5 mg Tamoxifen (Sigma, T5648; dissolved in medium chain fatty acids (Sigma, C8267)) at 10 mg/mL per day for 5 consecutive days to induce Cre activity. Both male and female mice between 8-12 weeks of age were used in all the studies. All mice were housed in the breeding core at the University of Wisconsin-Madison. All protocols were approved by the University of Wisconsin-Madison Institutional Animal Care and Use Committee.

### Flow cytometry of hematopoietic populations

Femurs and tibias were harvested from 8-12 week-old mice and crushed in phosphate-buffered saline (PBS, Corning, 21-040-CV) containing 2% heat inactivated fetal bovine serum with mortar and pestle. PB was collected by cardiac puncture. RBCs were removed by lysis with BD PharmLyse Lysing Buffer (BD Biosciences, 555899). For HSPC isolation, BM cells were pre-enriched for c-Kit+ cells using c-Kit microBeads (Miltenyi Biotec, 130-091-224) and a Midi MACS separator (Miltenyi Biotec, 130-042-301). BM, PB, or c-Kit-enriched cells were then stained for 30 min at 4℃ with combinations of antibodies to the following cell surface markers conjugated with different fluorochromes: CD3 (BioLegend, 17A2), CD11b (BioLegend, M1/70), Ter119 (BioLegend, TER119), Gr1 (BioLegend, RB6-8C5), B220 (BioLegend, RA3-6B2), c-Kit (BioLegend, 2B8), Sca-1 (BioLegend, D7), CD127 (BioLegend, SB/199), CD135 (BioLegend, A2F10). HSPC isolation was performed on BD FACSAria and BD FACSDiscover S8 Cell sorters. Data collection was performed on BD LSRII Fortessa and data analysis was performed using FlowJo v10.10.

### Mass spectrometric detection of GABA

For LC/MS/MS analysis of GABA, mouse bones were prepared by homogenization, precipitation, and filtration. Details are provided in Supplementary Methods.

### In vitro B cell differentiation assay

OP9 stromal cells were purchased from American Type Culture Collection (CRL-2749) and were grown as previously described (Shao *et al*., 2021). For each experiment, we harvested BM from 1 or 2 WT or Gad1/2 KO mice and sorted 4,000 LSK HSPCs into each well of a 12-well plate; replicates are 3 separate wells for each condition. Each well had a monolayer of OP9 cells at 70-80% confluency and 1 ml DMEM medium containing 10% FBS, 5 ng/ml Flt-3L (PeproTech, 250-31L) and 1 ng/ml IL-7 (R&D Systems, 407-ML-005). Experiments were repeated and the results of one representative experiment are shown. After 7 days of co-culturing, hematopoietic cells were mechanically detached, filtered and transferred to fresh OP9 monolayers. Cells were harvested 12 days after initial culturing and stained with Cd45 (BioLegend, 30-F11), B220 (BioLegend, RA3-6B2), Cd19 (BioLegend, 6D5) and Cd11b (Biolegend, M1/70) antibodies. Cells were analyzed by brightfield microscopy Nikon ECLIPSE TS2 and flow cytometry. Conditioned media was analyzed by mass spectrometry. Cell counts were determined by hemacytometer. For time course experiments, cells were harvested and analyzed at 3-day intervals after initiation of co-cultures. Flow cytometry analysis was performed using BD LSRII Fortessa and FlowJo v9/10.10.

### Bulk RNA-seq library preparation and sequencing

LSK cells from 8-12 week-old wild-type or *Cd19-Cre;Gad1/2^fl/fl^*, *Cdh5-CreERT2;Gad1/2^fl/fl^*, and *Tie2-Cre;Gad1/2^fl/fl^* models (triplicates for each group) were sorted using BD FACSAria Cell Sorter or BD FACSDiscover S8 Cell Sorter and collected in Trizol LS (Invitrogen). Nuclease-free water was added to adjust the volume to a final ratio of 3:1 for Trizol LS and sample. Samples were stored at -80℃ until ready for RNA extraction. Samples were thawed, equilibrated to room temperature, and 100 uL nuclease-free water was added to each sample to reduce the density of aqueous phase. 5PRIME Phase Lock Gel tubes (Quantabio) were used following the manufacturer instructions to separate the aqueous phase. Once the aqueous phase was separated, 25 ug of GenElute-LPA (Sigma, Cat.56575) was added to each sample for RNA precipitation. After adding isopropanol, samples were stored overnight at -20℃ for RNA precipitation. The pellet was then washed three times with 75% ethanol, resuspended in 10 uL nuclease-free water, and stored at -80℃ until library preparation.

RNA samples were quantified, and integrity was examined using Agilent 2100 Bioanalyzer RNA PicoChip. RNA libraries were prepared by UW-Madison Gene Expression Center using Takara SMARTer Stranded RNA v2 Prep Kit - Pico Input Mammalian (Takara Bio). RNA input for library preparation was 1-2.5 ng. The quality of libraries was examined using Agilent 4200 TapeStation. Sequencing was run on an Illumina NovaSeq X Plus single 2x150 bp lane at the UW-Madison Next Generation Sequencing Core.

### Bulk RNA-Seq analysis

Raw reads were checked for quality (FastQC v0.12.1) and adapter sequences were trimmed. Reads were mapped to the mouse reference genome GRCm39 using STAR aligner and read counts were normalized using RSEM by the UW-Madison Bioinformatics Resource Core. Low expression genes were filtered, library size was normalized and DGE (Differential Gene Expression) analysis was performed using EdgeR (v4.4.2). Principal component analysis identified an outlier that was removed from further analysis. The final analysis included n=3 samples of sorted HSPCs from WT, *Tie2-Cre;Gad1/2^fl/fl^*, and *Cdh5-CreERT2;Gad1/2^fl/fl^* BM samples, and n=2 from *Cd19-Cre;Gad1/2^fl/fl^*. Pairwise comparisons between WT samples and each mutant were performed using the quasi-likelihood F-test and genes with Log2 Fold Change > 1 and False Discovery Rate (FDR) < 0.1 were selected as DE genes. For heatmaps depicting unsupervised hierarchical clustering, genes that were differentially expressed in at least one mutant were used with the following parameters. GSEA (Gene Set Enrichment Analysis) of HSPCs isolated from mutants was performed using clusterProfiler (release 3.21) with GseaPreranked method. MSigDB Gene sets were downloaded using R package msigdbr (v7.5.1). Gene sets used for GSEA included HALLMARK and GO:BP sets. Rank scores for each pairwise comparison were created by multiplying the negative log of the p-value by the sign of the log fold change and ranking in descending order. A cutoff of adjusted p value < 0.05 was applied for GSEA and results were further filtered using the following criteria: adjusted p-value < 0.01, setSize >= 15 and <= 301, abs(NES) >= 2. Data was submitted to NCBI GEO under the accession number GSE294951.

### Statistics

The data was presented as mean ± sd. For the comparison between single experimental and control groups, Welch’s t test was used. For comparison between multiple groups, Analysis of Variance (ANOVA) was used, and Tukey’s HSD was used as the post-hoc test to determine differences between groups. P < 0.05 is considered as statistical significance. GraphPad Prism 10.4 was used for all analyses.

## Supporting information

Supplementary Table 1 DEG

Supplementary Table 2 GSEA

## Acknowledgments

The authors thank the University of Wisconsin Carbone Cancer Center Flow Cytometry Laboratory, supported by P30 CA014520, for use of its facilities and services, especially Kathryn Fox and Zach Stenerson for their technical assistance with panel optimization; Kathy Krentz at University of Wisconsin-Madison Animal Models Core for mouse model rederivation and cryopreservation; Megan Latsch and Jody Peter of the University of Wisconsin Biomedical Research Model Services for mouse colony management; Lauren Wells and Sandra Splinter BonDurant at University of Wisconsin-Madison Biotechnology Gene Expression Center (Research Resource Identifier – RRID:SCR_017757) for bulk RNA-sequencing library preparation; University of Wisconsin-Madison Next Gen DNA Sequencing Core for Illumina sequencing; Mark E. Berres at University of Wisconsin-Madison Biotechnology Center Bioinformatics Core Facility (Research Resource Identifier – RRID:SCR_017799) for sequencing reads alignment. Flow cytometry analyses and cell sorting utilized instruments that were supported by the following grants: BD LSR Fortessa (1S10OD018202-01), BD FACS AriaII BSL-2 Cell Sorter, “Jill” (1S10RR025483-01). Research reported in this publication was supported by the Department of Cell and Regenerative Biology at the University of Wisconsin-Madison.

## Author contributions

Conceptualization, O.J.T.; Data curation, C.D.; Formal Analysis, C.D., A.E.-B., M.P.H.; Funding Acquisition, O.J.T.; Investigation, C.D., A.E.-B., A.T.S., R.M.C., C.J.O., M.P.H., C.O.S.; Methodology, M.P.H., C.O.S.; Project Administration, O.J.T.; Resources, O.J.T.; Software, C.D.; Supervision, O.J.T.; Validation, C.D.; Visualization, C.D. and A.E.-B.; Writing - Original draft, C.D.; Writing – reviewing and editing, O.J.T..

## Supplementary Methods

### Mass spectrometric detection of GABA

All solvents for processing and mass spectometry analysis were Optima LCMS grade from Fischer Scientific (Waltham, MA), unless otherwise noted. Individual bones were weighed and placed in a 2-ml screwcap tube pre-loaded with 2.8mm ceramic beads (Fischer Scientific, Pittsburgh PA). Ice cold 150 mM ammonium bicarbonate (Sigma Aldrich, St. Louis MO) was added (5 mls/gram of tissue) and samples were placed on ice. Samples were then homogenized in an Omni 48-place bead mill homogenizer (Omni International, Kennesaw, GA) set to 6m/sec for 30 sec. Sample homogenization was repeated twice more to fully disrupt bone tissue placing samples on ice between replicates. Following processing 50 µl of bone homogenate was transferred to a 1.7 ml microcentrifuge tube and precipitated with 200 µl ACN/1% formic acid containing internal standard, d_6_-g-aminobutyric acid, (CDN resource laboratories Langley, BC). To remove insoluble material the samples were spun in a refrigerated centrifuge at 20,000 xg for 15 mins. The supernatant was filtered in a prewashed 96-well format Sirocco plate (Waters Corp. Milford, MA) into a 1-ml receiver plate according to the manufacturer’s protocol. The plate was then dried under nitrogen. Samples were resuspended in 150 ul 1% formic acid for LC/MS/MS analysis. Calibrators (0.75 ng/ml-500 ng/ml) and QC samples for g-aminobutyric acid (Sigma Aldrich, St. Louis MO) were prepared in water, as GABA free bone matrix is not available. The calibrators and QC samples were processed in tandem with unknown samples (addition of ISTD, precipitation and filtration).

For quantitative analysis samples, QCs and calibrators were injected in random order onto a 150 mM Intrada Amino Acid Column (Imtakt, Portland OR) using a Waters Acquity I-Class UPLC system (Waters, Milford MA). The column was held at 60 °C and the flow-rate was 0.500 ml/min.

Analytes were eluted from the column using a gradient. Solvent A was water/0.1% FA and solvent B was ACN/0.1% FA. Briefly, 7.5 µl of sample was injected at 30% B and the %B was ramped to 75% B in 5.5 minutes, then to 99% in 0.2 min. The %B was held at 99% for 1.3 minutes before returning to 30% in 0.2 min followed by a re-equilibration hold at 1.2 minutes before injection of the next samples. Eluate from the column was analyzed in positive ion mode using a QTrap 5500 hybrid triple quadrupole mass spectrometer (SCIEX, Framingham MA) operating in multiple-reaction-mode (MRM) under conditions optimized for detection of the analyte and internal standard. For g-aminobutyric acid the transitions were: parent ion 104.05/product ions 87, 86 and 69. The transitions for d_6_-g-aminobutyric acid: parent 110.09/product ions 93, 92.5 and 73.10. All transitions had a 50 msec dwell time. Triplicate injections of samples, calibrators and QCs were used for quantitative analysis allowing calculation of mean and standard deviation, blanks were run between each injection. The mean area under the curve (AUC) of the analyte relative to ISTD was used to construct a quadratic fit for the calibration curve in the MultiQuant software (SCIEX, Framingham, MA). Calibrators were excluded from quantitation models if their calculated concentrations were >15% different from theoretical. For all calibrators, samples, or QCs if the calculated %RSD of the triplicate injections was >15% samples were not considered valid.

All calibration curves had r-values ≥ 0.995. For all assays QC samples at each concentration fell within 15% of theoretical concentrations. The lower limit of detection was based on a signal to noise value of three (determined in MultiQuant). The lower limit of quantitation for each assay was based on either a signal to noise value of ten, or set at a g-aminobutyric acid concentration equal to the lowest concentration calibrator, whichever was higher. The upper limit of quantitation for each assay was set at the highest calibrator meeting the ±15% of theoretical metric.

---

**Supplementary Figure 1.**
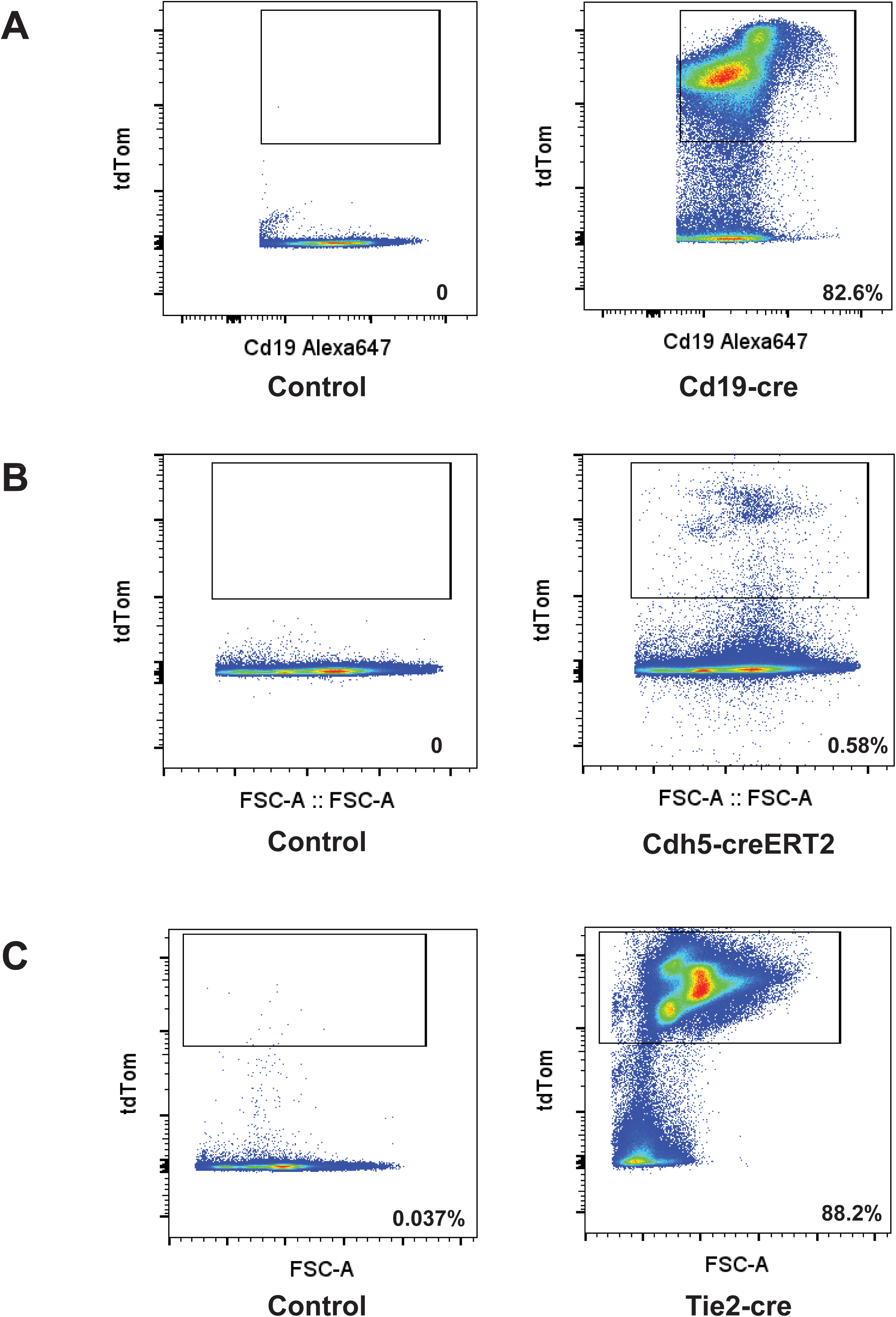
Genetic knockout efficiency in Gad1/2^fl/fl^ mutants. (A) Gad1 genetic knockout efficiency shown by the percentage of tdTomato+ CD19+ cells in*Cd19-Cre;Gad1/2^fl/fl^;Rosa26^tdTomato^* mice. (B) Gad1 genetic knockout efficiency shown by the presence of tdTomato+ cells in the whole BM cells in *Cdh5-CreERT2;Gad1/2^fl/fl^;Rosa26^tdTomato^* mice. (C) Gad1 genetic knockout efficiency shown by the percentage of tdTomato+ cells in the whole BM cells in *Tie2-Cre;Gad1/2^fl/fl^;Rosa26^tdTomato^*mice.

**Supplementary Figure 2.**
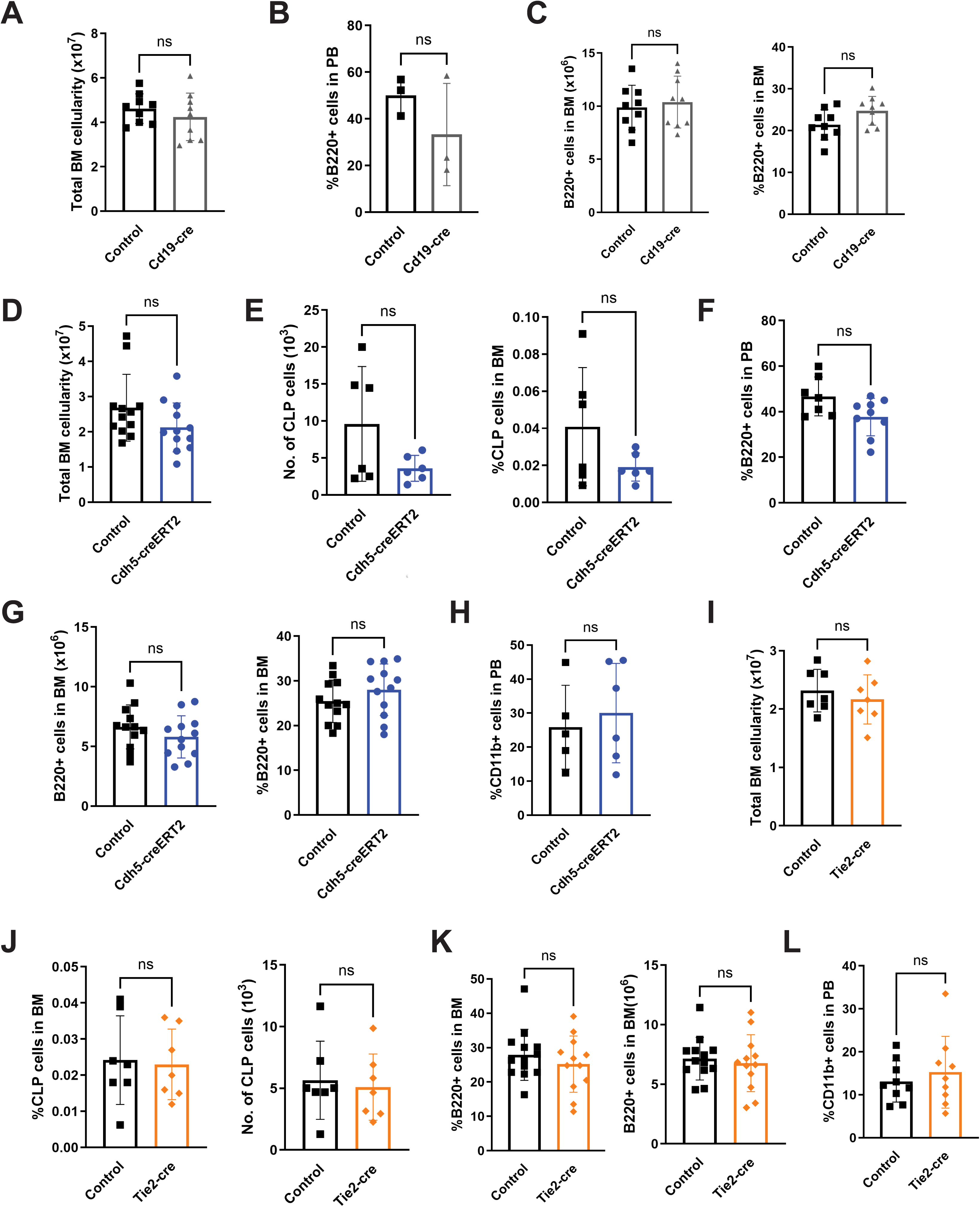
Hematopoietic profiles of Gad1/2^fl/fl^ mutants. (A) Total BM cells from 1 femur and 1 tibia in *Cd19-Cre;Gad1/2^fl/fl^;Rosa26^tdTomato^* and control mice (n = 9 mutants vs n = 9 control). (B) Quantification of B220+ cell percentage in the PB in *Cd19-Cre;Gad1/2^fl/fl^;Rosa26^tdTomato^* mice and control mice (n = 3 mutants vs n = 3 control). (C) Quantification of BM B220+ cells in *Cd19-Cre;Gad1/2^fl/fl^;Rosa26^tdTomato^* mice and control mice (n = 9 mutants vs n = 9 control). (D) Total BM cells from 1 femur and 1 tibia in *Cdh5-CreERT2;Gad1/2^fl/fl^;Rosa26^tdTomato^* mice and control mice (n = 12 mutants vs n = 12 control). (E) Quantification of common lymphoid progenitors (CLPs; Lin-/Sca-1-lo/c-Kit-lo/CD127+/CD135+) in *Cdh5-CreERT2;Gad1/2^fl/fl^;Rosa26^tdTomato^* and control mice (n = 6 mutants vs n = 6 control). (F) Percentage of B220+ cells in the PB in *Cdh5-CreERT2;Gad1/2^fl/fl^;Rosa26^tdTomato^* mice and control mice (n = 9 mutants vs n = 7 controls). (G) Quantification of BM B220+ cell in *Cdh5-CreERT2;Gad1/2^fl/fl^;Rosa26^tdTomato^* mice and control mice (n = 12 mutants vs n = 12 control). (H) Percentage of CD11b+ cells in the PB in *Cdh5-CreERT2;Gad1/2^fl/fl^;Rosa26^tdTomato^* and control mice (n = 6 mutants vs n = 5 control). (I) Total BM cells from 1 femur and 1 tibia in *Tie2-Cre;Gad1/2^fl/fl^;Rosa26^tdTomato^* mice and control mice (n = 7 mutants vs n = 7 control). (J) Quantification of CLPs (Lin-/Sca-1-lo/c-Kit-lo/CD127+/CD135+) in *Tie2-Cre;Gad1/2^fl/fl^;Rosa26^tdTomato^*and control mice (n = 7 mutants vs n = 7 control). (K) Quantification of BM B cell abundance in *Tie2-Cre;Gad1/2^fl/fl^;Rosa26^tdTomato^* mice and control mice (n = 12 mutants vs n = 12 control). (L) Percentage of CD11b+ cells in the PB in *Tie2-Cre;Gad1/2^fl/fl^;Rosa26^tdTomato^* and control mice (n = 9 mutants vs n = 9 control). Data is represented as mean ± SEM (*p<0.05), Welch’s test was performed.

**Supplementary Figure 3.**
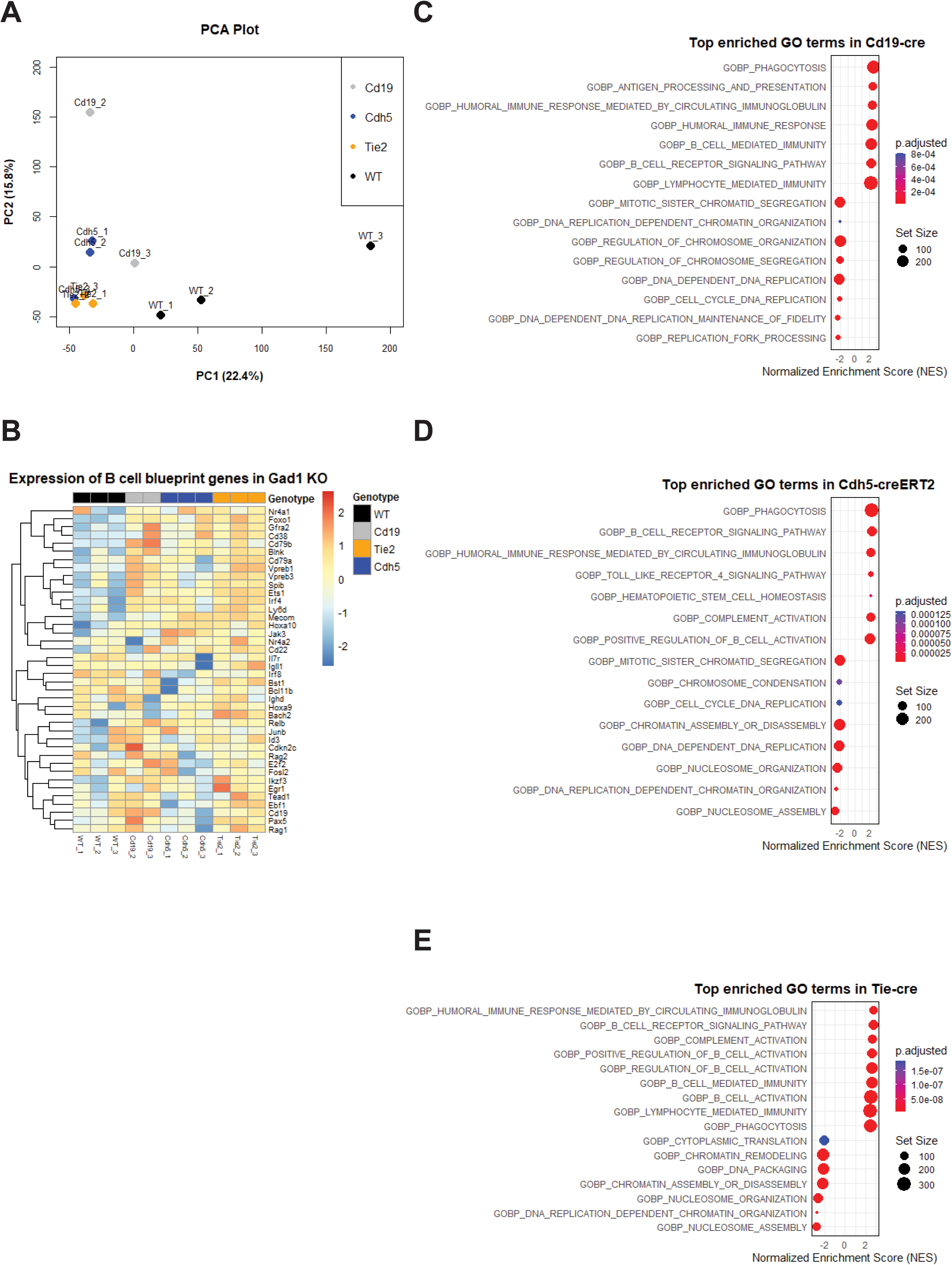
Gene expression profiles of Gad1/2^fl/fl^ mutants. (A) Distinct gene expression profiles of WT and *Gad1/2^fl/fl^* mutant models shown by PCA plot. (B) Heatmap of normalized expression levels of B cell blueprint genes in *Gad1/2^fl/fl^* mutant models. (C) GSEA result showing top enriched GO terms in *Cd19-Cre;Gad1/2^fl/fl^;Rosa26^tdTomato^* mutants. (D) GSEA result showing top enriched GO terms in *Cdh5-CreERT2;Gad1/2^fl/fl^;Rosa26^tdTomato^* mutants. (E) GSEA result showing top enriched GO terms from *Tie2-Cre;Gad1/2^fl/fl^;Rosa26^tdTomato^* mutants.

